# MMP9-Mediated Junctional Disassembly and Golgi Apparatus Disruption Compromise Astrocytic Integrity upon Human Coronavirus OC43 Infection

**DOI:** 10.64898/2026.11.19.767860

**Authors:** Souvik Karmakar, Jayasri Das Sarma

## Abstract

Neuropathogenesis of human coronavirus OC43 (HCoV-OC43) involves astrocyte infection, but it is unknown whether it disrupts junctional proteins essential for CNS homeostasis and blood-brain barrier integrity. We demonstrate that HCoV-OC43 infection of primary mouse astrocytes broadly disrupts junctional architecture, reducing expression and altering the localization of gap junction (Cx43), tight junction (Occludin, ZO-1), and adherens junction (β-catenin, E-cadherin) proteins. This disassembly coincides with Golgi apparatus fragmentation, implicating trafficking defects and NF-κB-driven upregulation of MMP9, accompanied by elevated levels of inflammatory cytokines, chemokines, and interferons. ROS/ERK-MAPK signaling does not contribute to MMP9 induction. Critically, pharmacological MMP inhibition restores junctional protein expression and localization, establishing MMP9 as a functional mediator of junctional disruption. These findings reveal that HCoV-OC43 compromises astrocytic junctional integrity through NF-κB-mediated MMP9 proteolysis and Golgi trafficking dysfunction, thereby identifying potential therapeutic targets for coronavirus-associated neurological disease.

**Importance:** Astrocytes play an important role in maintaining central nervous system homeostasis, yet how coronavirus infection affects their cellular integrity remains poorly understood. This study demonstrates that HCoV-OC43 infection disrupts multiple junctional complexes in primary mouse astrocytes through mechanisms involving Golgi apparatus disruption and an NF-κB-associated inflammatory response accompanied by increased matrix metalloproteinase 9 expression. These changes may compromise junctional protein trafficking and promote their alteration in expression and localization. Importantly, pharmacological inhibition of matrix metalloproteinase activity restores the expression and membrane localization of junctional proteins, highlighting a functional contribution of proteolytic activity to this process. These findings provide insight into how coronavirus infection can simultaneously alter astrocytic structural organization and inflammatory responses. Current findings also identify matrix metalloproteinase activity as a potential therapeutic target for preserving astrocytic integrity during coronavirus-associated central nervous system infections.

## Introduction

Coronaviruses, including the highly pathogenic SARS-CoV, MERS-CoV, and SARS-CoV-2, as well as endemic seasonal strains OC43, 229E, NL63, and HKU1, are established respiratory pathogens with recognized neurotropic potential^1^. These viruses can access the CNS through olfactory, hematogenous, and neural routes, and clinical and experimental evidence increasingly implicates coronavirus neuroinfection in neurological manifestations and cognitive impairment^2^. Within the CNS, astrocytes are essential regulators of homeostasis and blood-brain barrier (BBB) integrity. Through perivascular endfeet that ensheathe cerebral vasculature, astrocytes maintain the neurovascular unit and regulate endothelial barrier properties via junctional protein networks^3^. These networks comprise three functionally distinct classes: gap junctions, primarily Connexin 43 (Cx43), which mediate direct intercellular communication and metabolite exchange between astrocytes and between astrocytes and oligodendrocytes; tight junctions, including Occludin and ZO-1, which establish paracellular barrier integrity at the BBB interface; and adherens junctions, including β-catenin and E-cadherin, which maintain cell-cell adhesion, cytoskeletal linkage, and tissue organization^4,5^. Disruption of these junctional complexes has been linked to neuroinflammation, BBB compromise, and neurodegeneration. Viral pathogens manipulate junctional proteins to cross tissue barriers, facilitate intercellular spread, and evade immune surveillance^6–8^. However, whether coronavirus infection broadly remodels astrocytic junctional architecture beyond gap junctions remains poorly defined.

We previously established that β-coronaviruses selectively disrupt astrocytic gap junction communication. The murine β-coronavirus MHV-A59 downregulates Cx43 through ER/ERGIC retention and impaired trafficking, disrupting functional gap junction intercellular communication (GJIC) in both primary astrocytes and the infected mouse brain^9,10^. *In vivo*, this acute Cx43 loss is accompanied by persistent downregulation of the oligodendroglial gap junction protein Cx47^10^. Cx43 forms heterotypic gap junctions with oligodendrocytic Cx47 to support metabolic coupling essential for myelin maintenance; thus, sustained Cx47 depletion correlates with loss of the myelin marker proteolipid protein and progressive chronic demyelination^10,11^. In lung epithelial cells, HCoV-OC43 similarly reduces Cx43, traps Cx43 in the ERGIC region, disrupts Golgi-dependent trafficking, and elevates ER stress, impairing both GJIC and hemichannel formation^12^. Mechanistically, MHV-A59-induced ER stress downregulates the ER-resident molecular chaperone ERp29, which governs Cx43 oligomerization and surface delivery; restoring ERp29 via the chemical chaperone 4-phenylbutyric acid (4-PBA) rescues Cx43 and Cx47 expression and GJIC, restricts viral infectivity, and mitigates chronic demyelination *in vivo*^13,14^. While these studies established a conserved β-coronavirus mechanism of gap junction disruption linked to trafficking defects and ER stress, they examined only gap junction proteins. Astrocytes also depend on tight and adherens junctions for BBB support and intercellular integrity; therefore, determining whether β-coronavirus infection disrupts these junctional classes and identifying the cellular processes that drive such broad remodeling are essential next steps.

In this study, we demonstrate that HCoV-OC43 infection of primary mouse astrocytes broadly dismantles junctional architecture beyond gap junctions, reducing expression and mislocalizing tight junction (Occludin, ZO-1) and adherens junction (β-catenin, E-cadherin) proteins in addition to Cx43. This junctional disassembly coincides with fragmentation of the Golgi apparatus, extending our prior observation of coronavirus-induced Golgi disruption from epithelial cells to astrocytes, and with NF-κB-driven MMP9 upregulation alongside elevated inflammatory cytokines, chemokines, and interferons; ROS/ERK-MAPK signaling does not contribute to MMP9 induction under these conditions. Critically, pharmacological MMP inhibition with GM6001 restores junctional protein expression and localization across all three junctional classes, establishing MMP9 as a functional mediator of junctional disruption and revealing a proteolytic mechanism distinct from the ERp29-dependent trafficking pathway previously described. These findings reveal that HCoV-OC43 compromises astrocytic junctional integrity through NF-κB-mediated MMP9 proteolysis and Golgi trafficking dysfunction, a mechanism directly relevant to neurological complications in SARS-CoV-2 infection, identifying astrocytic MMP9 as a therapeutic target for coronavirus-induced CNS disease.

### STAR Methods

### Key Resources Table

| Reagent and Virus Strains | Source | Identifier |
|---|---|---|
| HCoV-OC43 | BEI Resources, NIAID, NIH | Cat# NR-56241 |
| **Chemicals, Peptides, and Recombinant Proteins** |  |  |
| Trypsin (10 mg/mL) | Sigma-Aldrich | Cat# T9201 |
| DNase I (0.1 mg/mL) | Sigma-Aldrich | Cat# D5025 |
| DMEM | Gibco | Cat# 12100-046 |
| FBS | Gibco | Cat#10082-147 |
| L-Glutamine (200 mM) | Gibco | Cat# 25030-081 |
| Non-Essential Amino Acids (100×) | Gibco | Cat# M7145 |
| Penicillin-Streptomycin | Gibco | Cat# 15140-122 |
| HBSS | Gibco | Cat# 14185-052 |
| 0.25% Trypsin | Gibco | Cat# 15050057 |
| 0.5% Trypsin | Gibco | Cat# 15400054 |
| GM6001 (Ilomastat) | MedChemExpress | Cat# HY-15768 |
| DCFDA | Sigma-Aldrich | Cat# D6883 |
| MTT | SRL | Cat# 33611 |
| DMSO | Sigma-Aldrich | Cat# D2650 |
| PVDF membranes | Millipore | Cat# IPVH00010 |
| SuperSignal West Pico PLUS Chemiluminescent Substrate | Thermo Scientific | Cat# 34580 |
| TRIzol Reagent | Invitrogen | Cat# 15596026 |
| Pierce BCA Protein Assay Kit | Thermo Scientific | Cat# 23225 |
| Paraformaldehyde | SRL | Cat# 94464 |
| Triton X-100 | Sigma-Aldrich | Cat# T8787 |
| Mowiol Mounting Medium with DAPI | N/A | N/A |
| **Critical Commercial Assays** |  |  |
| High-Capacity cDNA Reverse Transcription Kit | Applied Biosystems | Cat# 4368814 |
| iTaq Universal SYBR Green Supermix | Bio-Rad | Cat# 1725124 |
| **Experimental Models: Organisms/Strains** |  |  |
| C57BL/6 mice (neonatal, P0-P1) | IISER Kolkata Animal Facility | N/A |
| Vero cells | Gift from Dr. Sourish Ghosh (Indian Institute of Chemical Biology, Kolkata, India) | N/A |
| **Antibodies** |  |  |
| Anti-HCoV-OC43 (monoclonal) | Gift from Dr. Brenda Hogue, Arizona State University | N/A |
| Anti-GFAP | Sigma-Aldrich | Cat# G9269 |
| Anti-Connexin 43 (IF) | Sigma-Aldrich | Cat# C6219 |
| Anti-Connexin 43 (WB) | Abclonal | Cat# A11752 |
| Anti-Occludin | Proteintech | Cat# 27260-1-AP |
| Anti-ZO-1 | Invitrogen | Cat# 61-7300 |
| Anti-β-catenin | Cell Signaling Technology | Cat# D10A8 |
| Anti-E-cadherin (IF) | Proteintech | Cat# 20874-1-AP |
| Anti-E-cadherin (WB) | Affinity Biosciences | Cat# AF7718 |
| Anti-TGN46 | Abclonal | Cat# A19618 |
| Anti-p65/RelA | Bio-Bharati Life Science | Cat# BB-AB0070 |
| Anti-IKKβ | Bio-Bharati Life Science | Cat# BB-AB0094 |
| Anti-ERK2 | Bio-Bharati Life Science | Cat# BB-AB0135 |
| Anti-β-actin | Invitrogen | Cat# MA1-140 |
| Alexa 488 Goat Anti-Rabbit | Invitrogen | Cat# A-11034 |
| Alexa 488 Goat Anti-Mouse | Invitrogen | Cat# A-11001 |
| Alexa 568 Donkey Anti-Rabbit | Invitrogen | Cat# A-10042 |
| Alexa 568 Donkey Anti-Mouse | Invitrogen | Cat# A-10037 |
| HRP-conjugated Rabbit Secondary | Invitrogen | Cat# 31460 |
| HRP-conjugated Mouse Secondary | Invitrogen | Cat# 31430 |
| **Oligonucleotides** |  |  |
| See Table S1 for primer sequences | N/A | N/A |
| **Software and Algorithms** |  |  |
| ImageJ | NIH | N/A |
| NIS-Elements ER | Nikon | N/A |
| FlowJo | BD Biosciences | N/A |
| GENESys | Syngene | N/A |
| CFX Maestro | Bio-Rad | N/A |
| GraphPad Prism 8 | GraphPad Software | N/A |
| STRING (v12) | STRING Consortium | N/A |

### Experimental Model and Subject Details Primary Astrocyte Culture

Primary astrocytes were isolated from neonatal C57BL/6 mice (postnatal day 0-1) as described previously^15,16^. Following cranium removal, brains were dissected, meninges carefully removed, and tissues mechanically dissociated. The cell suspension was enzymatically digested in HBSS (Gibco) containing 0.25 mL of 10 mg/mL trypsin (Sigma-Aldrich) and 0.25 mL of 0.1 mg/mL DNase I (Sigma-Aldrich) per brain for 30 min at 37 °C with rocking. Cells were triturated in the presence of 0.25 mL FBS per brain and pelleted at 600 × g for 10 min. The pellet was resuspended in HBSS, passed through a 70-μm nylon mesh, and centrifuged again (600 × g, 10 min). Cells were resuspended in Dulbecco’s Modified Eagle Medium (DMEM; Gibco) supplemented with 10% FBS (Gibco), 1% non-essential amino acids, 1% L-glutamine (Gibco), and 1% penicillin–streptomycin (Gibco). After 24 h, non-adherent cells were removed and replaced with astrocyte maintenance medium (DMEM, 10% FBS, 0.1% L-glutamine, 1% penicillin–streptomycin), with medium changes every 2-3 days. To enrich astrocytes, confluent mixed glial cultures were maintained without medium change for 7-10 days to promote differential adhesion. Microglia were removed by orbital shaking at 180 rpm for 45 min at 37 °C. The remaining adherent monolayer, enriched for astrocytes, was used for all subsequent experiments. Astrocyte purity was confirmed by immunostaining for glial fibrillary acidic protein (GFAP).

## Animals

All animal experiments were conducted in strict accordance with the guidelines of the Institutional Animal Ethics Committee (IAEC) of the Indian Institute of Science Education and Research (IISER) Kolkata, under the Committee for Control and Supervision of Experiments on Animals (CCSEA) of the Government of India. Experimental protocols for the use of neonatal C57BL/6 mice were approved under protocol number IISERK/IAEC/AP/2019/29.01, SL No. 43. The protocol was amended and extended in the year 2026 under protocol number IISERK/IAEC/AP/2026/172.

## Method Details

### Virus Propagation and Infection

HCoV-OC43 (BEI Resources, Cat# NR-56241) was propagated and titrated in Vero cells as described previously^12^. Primary astrocytes were seeded on etched coverslips or culture dishes and infected at a multiplicity of infection (MOI) of 1. After 2 h of adsorption at 37 °C, the inoculum was removed, and fresh culture medium was added. Mock-infected controls were treated identically without virus exposure.

## Immunofluorescence Staining

Mock-infected and HCoV-OC43-infected primary astrocytes were fixed at the indicated time points with 4% paraformaldehyde for 15 min at room temperature. Cells were permeabilized with 0.2% Triton X-100 in PBS for 30 min, blocked in 2.5% heat-inactivated goat serum in PBS containing 0.2% Triton X-100 for 30 min at room temperature, and incubated overnight at 4 °C with primary antibodies (*see Key Resources Table for dilutions*). After three washes with 0.2% Triton X-100 in PBS, coverslips were incubated with secondary antibodies (*see Key Resources Table*) for 75 min at room temperature, washed, and mounted in Mowiol mounting medium containing DAPI. The images were taken using a Nikon Eclipse Ti2 inverted microscope, equipped with the AXR confocal scanning module, controlled by NIS-Elements ER software, Japan. Images were processed with ImageJ software.

## Western Blotting

At 2 days post-infection, cells were rinsed twice with ice-cold PBS and lysed in RIPA buffer (50 mM Tris, 150 mM NaCl, 0.1% SDS, 0.5% sodium deoxycholate, 1% Triton X-100) supplemented with EDTA-free protease and phosphatase inhibitors (1 mM sodium orthovanadate, 10 mM sodium fluoride, 10 mM sodium pyrophosphate decahydrate). Lysis proceeded on ice for 90 min with intermittent vortexing every 15 min, followed by brief sonication. Lysates were clarified by centrifugation at 13,200 rpm for 20 min at 4 °C. Protein concentration was determined using the Pierce BCA Protein Assay Kit (Thermo Scientific). Equal amounts of total protein (30 μg) were resolved by 10% SDS-PAGE and transferred to PVDF membranes (Millipore). Membranes were blocked with 10% skim milk in TBS containing 0.1% Tween-20 (TBST) for 1 h at room temperature and probed overnight at 4 °C with primary antibodies (see Key Resources Table). After three TBST washes (10 min each), membranes were incubated with HRP-conjugated secondary antibodies (see Key Resources Table) for 1 h at room temperature. Protein bands were visualized using SuperSignal West Pico Chemiluminescent Substrate (Thermo Scientific) and imaged with a Syngene G: Box ChemiDoc system (GENESys software). Membranes were re-probed with anti-β-actin to confirm equal loading. Densitometric analysis was performed with ImageJ.

## Quantitative Real-Time PCR

Total RNA was extracted from mock-infected and HCoV-OC43-infected primary astrocytes using TRIzol reagent (Invitrogen). RNA concentration and purity were assessed with a NanoDrop ND-2000 spectrophotometer (Thermo Scientific). One microgram of total RNA per sample was reverse-transcribed using the High-Capacity cDNA Reverse Transcription Kit (Applied Biosystems). qPCR was performed with iTaq Universal SYBR Green Supermix (Bio-Rad) on a Bio-Rad CFX Real-Time PCR Detection System under the following conditions: 95 °C for 7 min, followed by 40 cycles of 95 °C for 10 s and 60 °C for 30 s, with melting curve analysis. Each reaction was performed in technical quadruplicate. Relative gene expression was calculated by the ΔΔCT method with GAPDH as the endogenous control and expressed as fold change relative to mock-infected controls. Primer sequences are provided in Table S1.

## ROS Measurement by DCFDA

Cellular ROS levels were assessed using 2′,7′-dichlorofluorescin diacetate (DCFDA) as described. At 2 days post-infection, primary astrocytes were washed with warm PBS and incubated with 7.5 μM DCFDA in serum-free DMEM for 30 min at 37 °C in 5% CO2. Cells were harvested, resuspended in PBS, and analyzed by flow cytometry (BD FACSVerse) in the FITC channel. Data were analyzed with FlowJo software, and ROS levels in infected cells were expressed relative to mock-infected controls.

## MMP Inhibition with GM6001

To assess the contribution of MMP activity to junctional protein remodeling, primary astrocytes were infected with HCoV-OC43 at an MOI of 1. Following adsorption and inoculum removal, fresh culture medium was added. At 1 day post-infection, infected cells were treated with the broad-spectrum MMP inhibitor GM6001 at a concentration of 25 μM. Untreated infected cells received the corresponding vehicle and served as infection controls. Cells were harvested at 2 days post-infection for immunofluorescence and immunoblot analyses.

## Cell Viability Assay

GM6001 cytotoxicity was assessed by MTT assay, following the protocol described previously^17,18^. Primary astrocytes were seeded in 96-well plates at 1 × 10 ^5^ cells/mL (100 μL/well) and cultured for 7 days, with medium changes every 3 days. Cells were treated with GM6001 (0-200 μM) in astrocyte medium for 24 h. MTT solution (0.5 mg/mL in complete medium; Sigma-Aldrich) was then added for 6 h at 37 °C. Formazan crystals were solubilized in 100 μL DMSO per well, and absorbance was measured at 590 nm (BioTek Epoch Microplate Spectrophotometer). Each concentration was evaluated in four technical replicates across three independent experiments. Cell viability was calculated relative to vehicle-treated controls. The dose-response curve was fitted by nonlinear regression (four-parameter logistic model) using GraphPad Prism 8.

## STRING Pathway Analysis

Protein-protein interaction networks were generated using the STRING database (version 12). A predefined set of proteins associated with ERK/MAPK signaling and ROS detoxification/antioxidant activity was entered into STRING to examine predicted/known protein-protein interactions. The interaction network was visualized with a minimum confidence score of 0.7 (high confidence). Functional clustering was applied to examine associations among genes related to ERK/MAPK signaling and ROS detoxification/antioxidant activity.

## Quantification and Statistical Analysis

All data are presented as mean ± SEM from three independent experiments. Statistical comparisons between mock-infected and HCoV-OC43-infected groups were performed using an unpaired two-tailed Student’s *t*-test with Welch’s correction. A *p*-value < 0.05 was considered statistically significant. All analyses and graphical representations were performed using GraphPad Prism 8 (GraphPad Software, Inc.).

## Results

### HCoV-OC43 infection broadly remodels junctional protein architecture in primary astrocytes

Following the establishment of HCoV-OC43 infection in primary astrocytes (Supplementary Figure 1), we examined whether infection alters the expression and cellular distribution of major astrocytic junctional proteins. Cx43 showed a characteristic punctate distribution at the cell periphery in control astrocytes (Fig. 1A and A1), whereas HCoV-OC43-infected cells displayed a significant reduction in peripheral punctate Cx43 staining (Fig. 1B and B1, and white arrows in B1). The altered Cx43 distribution was further evident in the additional immunofluorescence images shown in Supplementary Figure 2. Consistent with the immunofluorescence findings, western blot analysis showed a significant reduction in Cx43 protein levels, as confirmed by densitometric analysis (Fig. 1C-D), accompanied by a significant decrease in Cx43 transcript levels (Fig. 1E).

**Figure 1.**
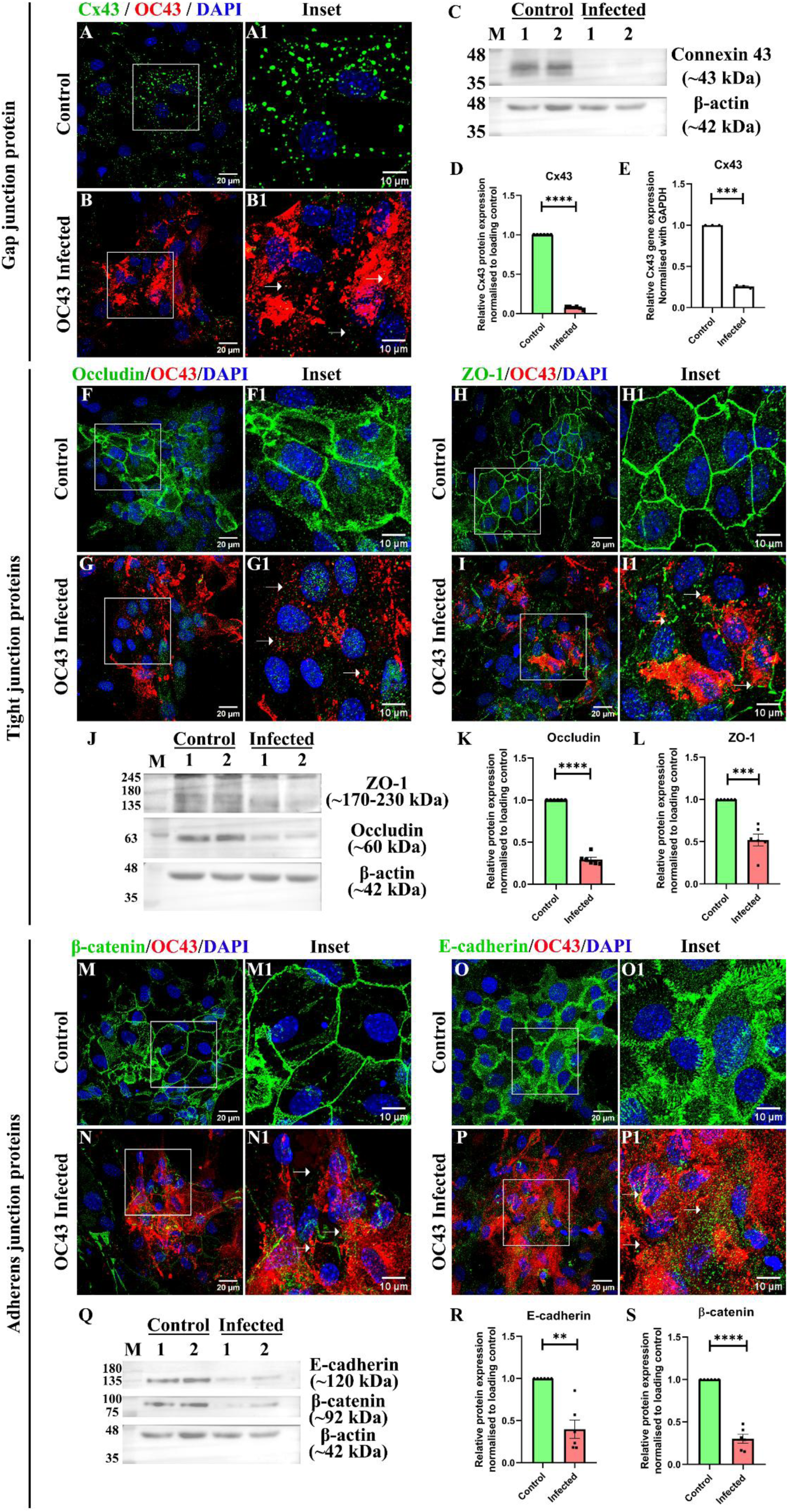
HCoV-OC43 infection broadly remodels junctional protein architecture in primary astrocytes. Primary astrocytes isolated from neonatal pups of C57BL/6 mice were infected with HCoV-OC43 at an MOI of 1 and analyzed at 2 days post-infection. (A-B) Control and HCoV-OC43-infected astrocytes were double-label immunostained for Cx43 (green), HCoV-OC43 (red), and counterstained with DAPI (blue). Control astrocytes showed characteristic punctate Cx43 staining at the cell periphery (A and A1), whereas HCoV-OC43-infected astrocytes displayed reduced peripheral Cx43 staining (B and B1, and white arrows in B1). (C) Representative immunoblot showing Cx43 and β-actin protein expression in control and HCoV-OC43-infected astrocytes. (D) Densitometric quantification of Cx43 protein expression normalized to β-actin. (E) Quantitative real-time PCR analysis of Cx43 transcript levels normalized to GAPDH. (F-G) Control and HCoV-OC43-infected astrocytes were double-label immunostained for Occludin (green), HCoV-OC43 (red), and DAPI (blue). Control astrocytes showed prominent peripheral Occludin localization (F and F1), whereas infected astrocytes displayed disrupted Occludin staining (G and G1, and white arrows in G1). (H-I) Control and HCoV-OC43-infected astrocytes were double-label immunostained for ZO-1 (green), HCoV-OC43 (red), and DAPI (blue). Control astrocytes showed prominent peripheral ZO-1 localization (H and H1), whereas infected astrocytes displayed disrupted ZO-1 staining (I and I1, and white arrows in I1). (J) Representative immunoblot showing ZO-1, Occludin, and β-actin protein expression in control and HCoV-OC43-infected astrocytes. (K-L) Densitometric quantification of Occludin and ZO-1 protein expression, respectively, normalized to β-actin. (M-N) Control and HCoV-OC43-infected astrocytes were double-label immunostained for β-catenin (green), HCoV-OC43 (red), and DAPI (blue). Control astrocytes showed prominent membrane-associated β-catenin localization (M and M1), whereas infected astrocytes displayed reduced and fragmented β-catenin staining (N and N1, and white arrows in N1). (O-P) Control and HCoV-OC43-infected astrocytes were immunostained for E-cadherin (green), HCoV-OC43 (red), and DAPI (blue). Control astrocytes showed membrane-associated E-cadherin localization (O and O1), whereas infected astrocytes displayed reduced and fragmented E-cadherin staining (P and P1, and white arrows in P1). (Q) Representative immunoblot showing E-cadherin, β-catenin, and β-actin protein expression in control and HCoV-OC43-infected astrocytes. (R-S) Densitometric quantification of E-cadherin and β-catenin protein expression, respectively, normalized to β-actin. Control and infected samples in the immunoblots represent technical replicates. Data are presented as mean ± SEM from three independent experiments. Statistical significance: **p < 0.01, ***p < 0.001, and ****p < 0.0001. Scale bars, 20 μm; inset scale bars, 10 μm.

We next examined the tight junction proteins Occludin and ZO-1. Both proteins showed prominent peripheral localization in control astrocytes (Fig. 1F and F1, and H and H1), whereas HCoV-OC43-infected astrocytes exhibited disrupted and diffuse staining patterns (Fig. 1G and G1, and I and I1, and white arrows in G1 and I1). These infection-associated changes in Occludin and ZO-1 localization were also observed in the additional immunofluorescence images presented in Supplementary Figure 3. Western blot analysis demonstrated significant reductions in Occludin and ZO-1 protein levels following infection, which were confirmed by densitometric quantification (Fig. 1J-L).

Adherens junctions also contribute to cell-cell adhesion and junctional organization; hence, we subsequently examined β-catenin and E-cadherin. Control astrocytes showed prominent membrane-associated localization of both proteins (Fig. 1M and M1, and O and O1), whereas HCoV-OC43-infected cells displayed reduced and fragmented membrane localization (Fig. 1N and N1, and P and P1, and white arrows in N1 and P1). Similar alterations in the distribution of β-catenin and E-cadherin were observed in the additional immunofluorescence images shown in Supplementary Figure 4. Western blot analysis further demonstrated significant reductions in β-catenin and E-cadherin protein levels, as confirmed by densitometric analysis (Fig. 1Q-S). Together, these findings demonstrate that HCoV-OC43 infection broadly remodels gap, tight, and adherens junctional proteins in primary astrocytes, affecting both their cellular localization and abundance.

## Disruption of the Golgi apparatus in murine primary astrocytes following HCoV-OC43 infection

The Golgi apparatus plays a pivotal role in post-translational modifications and protein trafficking. Since HCoV-OC43 infection was already found to disrupt the expression and localization of junctional proteins, the integrity of the Golgi apparatus was examined to assess whether alterations in host secretory pathways could be associated with the observed junctional changes. Immunofluorescence staining using TGN46, a marker of the trans-Golgi network, revealed intact perinuclear staining of the Golgi apparatus in control astrocytes (Fig. 2A-D). In contrast, HCoV-OC43-infected astrocytes at MOI 1 showed a significant loss of organized TGN46 staining, indicating disruption of Golgi organization (Fig. 2E-H, and white arrows in H). Western blot analysis further corroborated these observations, as it revealed reduced TGN46 protein expression in infected astrocytes compared to controls (Fig. 2I). Densitometric quantification confirmed a statistically significant decrease in protein levels (Fig. 2J). These findings demonstrate that HCoV-OC43 infection disrupts the Golgi apparatus and suggest that Golgi dysfunction may contribute to the altered organization of junctional proteins observed following infection.

**Figure 2.**
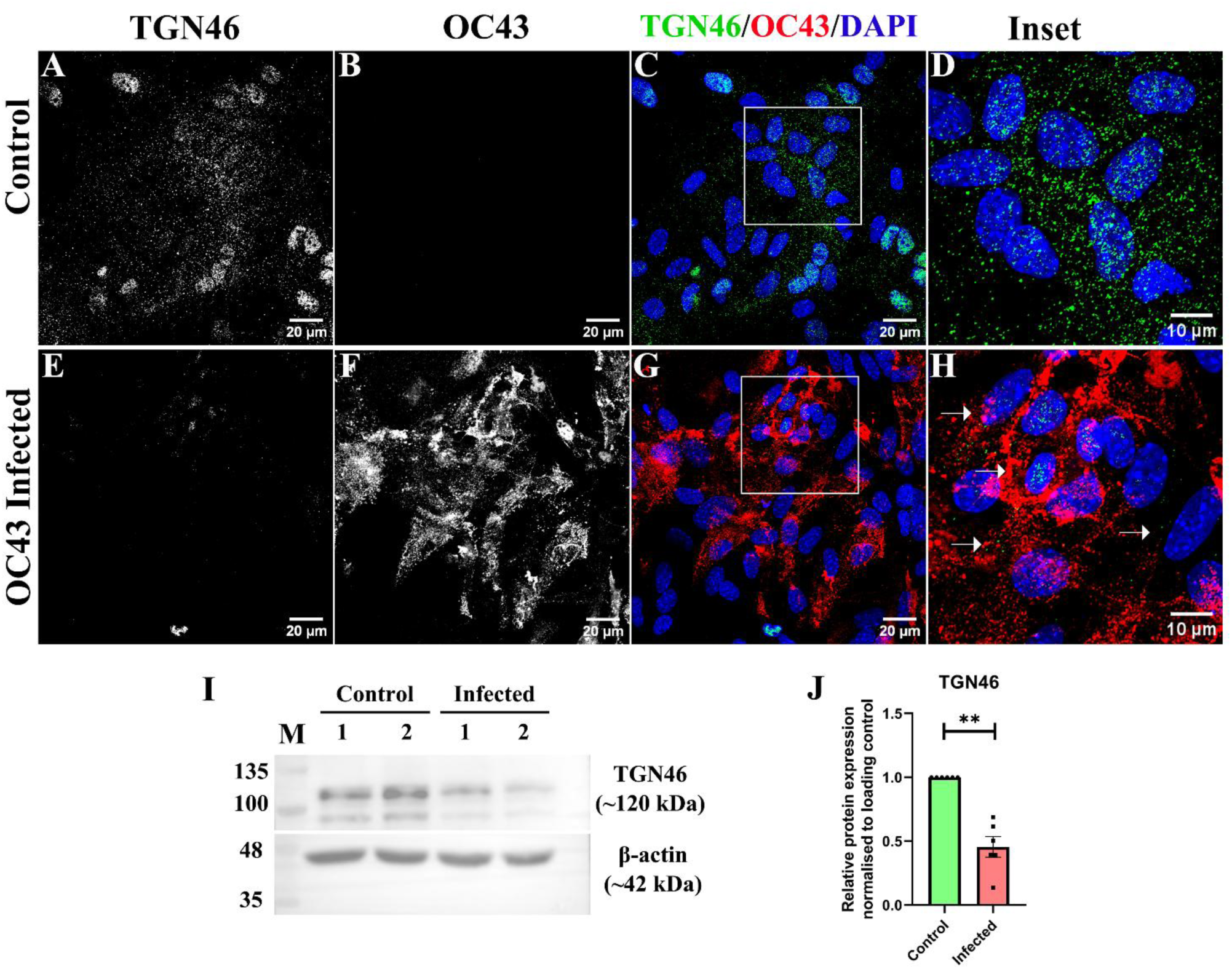
Disruption of the Golgi apparatus in primary astrocytes following HCoV-OC43 infection. Primary astrocytes were double-label immunostained with anti-HCoV-OC43 and anti-TGN46 antibodies and counterstained with DAPI. (A-D) Control astrocytes showed intact perinuclear Golgi organization as confirmed by the presence of TGN46 staining. (E-H) In contrast, infected astrocytes at MOI 1 exhibited reduced TGN46 staining and disruption of the perinuclear Golgi organization (white arrows, H). Panels C and G represent merged images. (I) Immunoblot analysis using total protein demonstrated reduced expression of TGN46 in infected astrocytes. (J) Densitometric quantification was performed using ImageJ. Control and infected lanes (1 and 2, respectively) represent technical replicates, and the M lane represents a pre-stained protein ladder. Data are shown as mean ± SEM from three independent experiments. Statistical significance: **p < 0.01. Scale bar: 20 µm; insets: 10 µm.

## HCoV-OC43 infection is associated with the activation of NF-κB signaling but excludes ROS/ERK-MAPK signaling pathway

We next examined potential cellular signaling pathways that could contribute to these infection-associated changes. Reactive oxygen species (ROS) are important signaling molecules involved in cellular responses to viral infection and have been implicated in the regulation of inflammatory and stress-associated pathways. We assessed ROS levels and the ERK/MAPK pathway activation. DCFDA-based flow cytometry revealed a significant reduction in ROS levels in HCoV-OC43-infected astrocytes compared with controls (Supplementary Figure 5A-C). We used STRING-based protein-protein interaction analysis of a predefined set of proteins associated with ROS/antioxidant processes and ERK/MAPK signaling. The resulting network showed interactions among proteins involved in both signaling pathways (Supplementary Figure 5D-E). However, immunoblot analysis showed reduced Erk2 protein expression following infection (Supplementary Figure 5F-G), providing no evidence of increased ERK/MAPK signaling upon HCoV-OC43 infection. These findings therefore do not support ROS/ERK-MAPK signaling as a major contributor to the cellular response associated with HCoV-OC43 infection.

As ROS/ERK-MAPK signaling did not appear to be increased following infection, we next examined the NF-κB pathway, a major regulator of innate and inflammatory responses to viral infection. Double-label immunofluorescence analysis showed increased p65/RelA staining in HCoV-OC43-infected astrocytes compared with control cells (Fig. 3A-H, and white arrows in H). Similarly, IKKβ staining was increased in infected astrocytes (Fig. 3I-P, and white arrows in P). These observations were supported by immunoblot analysis, which demonstrated increased p65/RelA and IKKβ protein levels in infected cells (Fig. 3Q-R), with densitometric analysis confirming significant increases in both proteins (Fig. 3S-T). At the transcript level, qRT-PCR showed significant upregulation of p65/RelA and IκBα transcripts following infection (Fig. 3U-V). Together, these findings demonstrate an HCoV-OC43-associated increase in NF-κB pathway components and inflammatory signaling in primary astrocytes. The subsequent induction of IκBα may reflect a compensatory response that may limit NF-κB signaling. Because NF-κB signaling can regulate inflammatory mediators and MMP expression, these findings raised the possibility that the NF-κB-associated response may contribute to the junctional remodeling observed following HCoV-OC43 infection. We therefore next examined MMP9 expression together with the associated cytokine and chemokine response.

**Figure 3:**
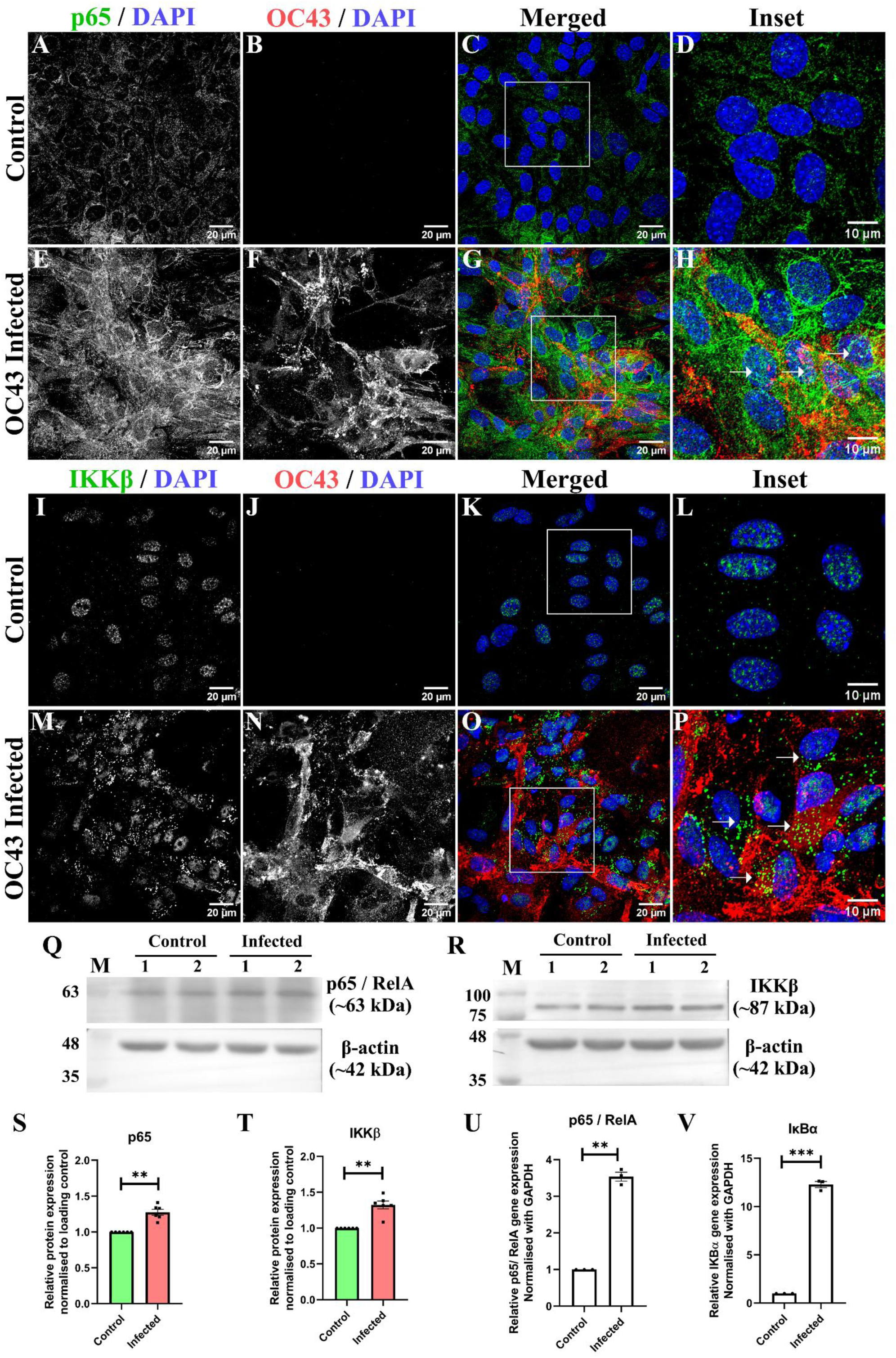
Activation of the NF-κB signaling pathway in primary astrocytes following HCoV-OC43 infection. Primary astrocytes were mock-infected or infected with HCoV-OC43 and analyzed for changes in NF-κB pathway-associated components. (A-H) Double-label immunofluorescence staining showed p65/RelA expression in control astrocytes and increased p65/RelA staining in HCoV-OC43-infected astrocytes. HCoV-OC43 staining confirmed infection, and nuclei were counterstained with DAPI. (H) The inset highlights increased p65/RelA staining in infected cells, with white arrows indicating representative regions. (I-P) Double-label immunofluorescence staining showed IKKβ expression in control astrocytes and increased IKKβ staining in HCoV-OC43-infected astrocytes compared with control cells. (P) The inset highlights increased IKKβ staining in infected cells, with white arrows indicating representative regions. (Q-R) Immunoblot analysis showed p65/RelA (∼63 kDa) and IKKβ (∼87 kDa) protein expression in control and HCoV-OC43-infected astrocytes. (S-T) Densitometric analysis using ImageJ showed significant changes in p65/RelA and IKKβ protein levels, respectively, in infected astrocytes compared with controls. (U-V) Quantitative real-time PCR analysis showed significant changes in p65/RelA and IκBα transcript levels, respectively, in HCoV-OC43-infected astrocytes compared with controls. Lanes 1 and 2 of the immunoblots represent technical replicates for control and infected samples, respectively, and lane M represents the pre-stained protein ladder. Data represent the mean ± SEM from three independent experiments. Statistical significance: *p < 0.05, **p < 0.01, and ***p < 0.001. Scale bar: 20 µm; insets: 10 µm.

## Induction of MMP9 expression in HCoV-OC43 infection in murine primary astrocytes

MMPs have been implicated in the remodeling of extracellular matrix components and junctional proteins. We therefore examined whether MMP-9 was induced following HCoV-OC43 infection. In HCoV-OC43-infected cells, a significant upregulation of MMP9 transcripts was observed with respect to controls (Fig. 4A). In addition, HCoV-OC43 infection induced a broad transcriptional inflammatory response. Pro-inflammatory cytokines and chemokines, including CCL5, IL-6, TNF-α, and IL-1β, were significantly upregulated in infected astrocytes compared with controls (Fig. 4B). Analysis of interferon responses showed significant upregulation of IFN-α and IFN-β transcripts, whereas IFN-γ expression remained unchanged (Fig. 4C). Interestingly, the anti-inflammatory cytokines IL-10 and TGF-β were also significantly elevated in infected cells (Fig. 4D). Together, these findings demonstrate that HCoV-OC43 infection is associated with MMP-9 induction and a broad inflammatory transcriptional response in primary astrocytes, providing another cellular response associated with the observed junctional remodeling.

**Figure 4:**
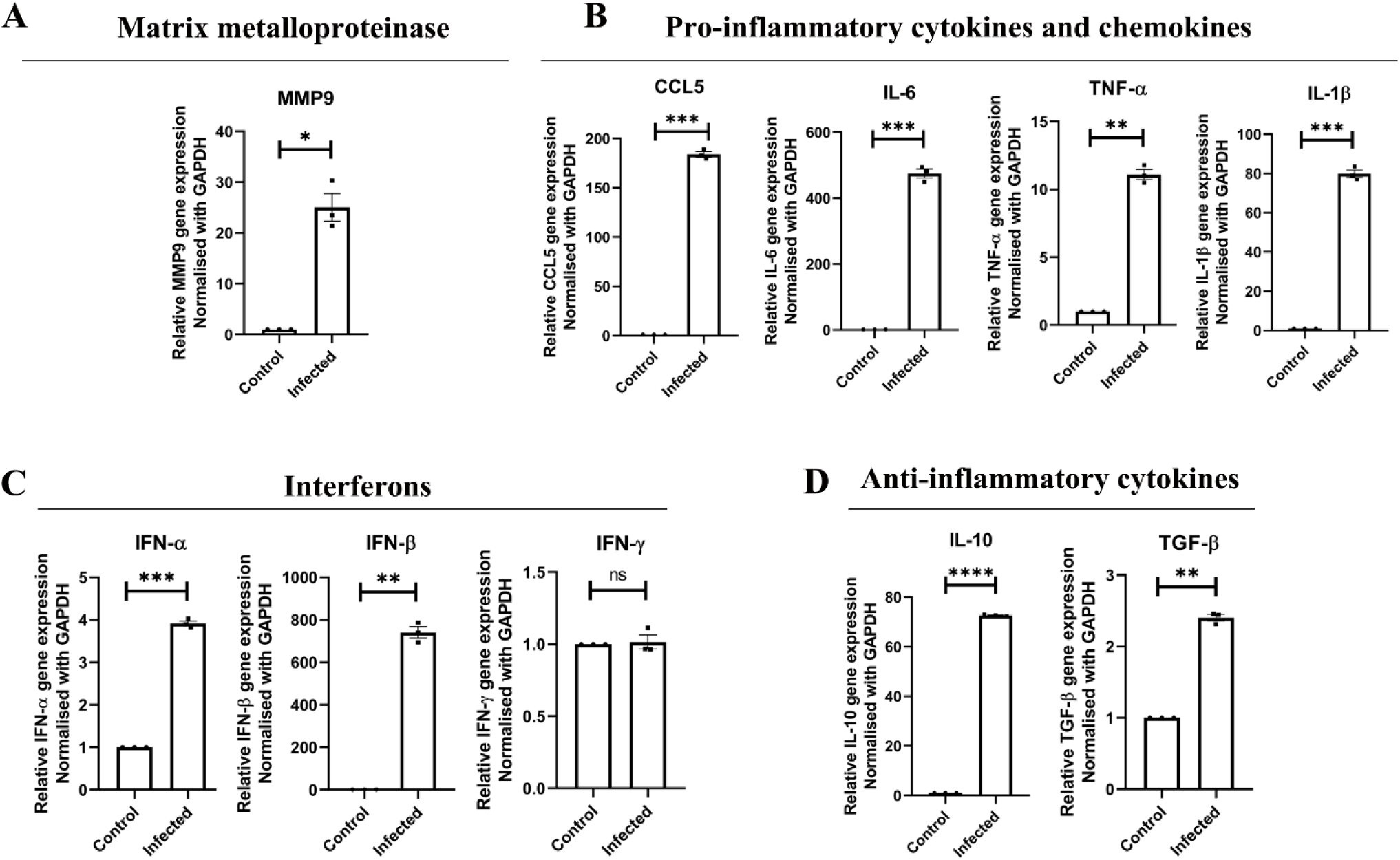
Differential expression of matrix metalloproteinase 9, cytokines, chemokines, and interferons in primary astrocytes following HCoV-OC43 infection. Primary astrocytes isolated from neonatal pups of C57BL/6 mice were infected with HCoV-OC43 at an MOI of 1, and transcript levels were analyzed by quantitative real-time PCR. (A) Infected astrocytes showed a significant increase in MMP9 expression. (B) Pro-inflammatory cytokines and chemokines showed strong induction, including CCL5, IL-6, TNF-α, and IL-1β, in infected astrocytes compared to controls. (C) Interferon responses revealed significant upregulation of IFN-α and IFN-β transcripts, while IFN-γ expression remained unchanged. (D) Anti-inflammatory cytokines IL-10 and TGF-β were also significantly elevated in infected cells. Gene expression levels were normalized to GAPDH. Data represent the mean ± SEM from three independent experiments. Statistical significance: *p < 0.05, **p < 0.01, ***p < 0.001, ****p < 0.0001, and ns = not significant.

## Determination of the working concentration of GM6001 for MMP inhibition in HCoV-OC43-infected primary astrocytes

To determine a suitable concentration of the broad-spectrum MMP inhibitor GM6001 for subsequent experiments, primary astrocytes were treated with increasing concentrations of GM6001 (0-200 µM), and cell viability was assessed using the MTT assay (Fig. 5A-B). No significant reduction in MTT-based cell viability was observed across the concentrations tested. Based on these findings, 25 µM GM6001 was selected as the working concentration for subsequent MMP inhibition experiments. The experimental treatment schedule consisted of HCoV-OC43 infection on Day 0, GM6001 treatment on Day 1, and sample collection on Day 2 post-infection (Fig. 5C). DIC imaging of HCoV-OC43-infected astrocytes without treatment and following GM6001 treatment showed no obvious treatment-associated cytotoxicity (Fig. 5D-E).

**Figure 5.**
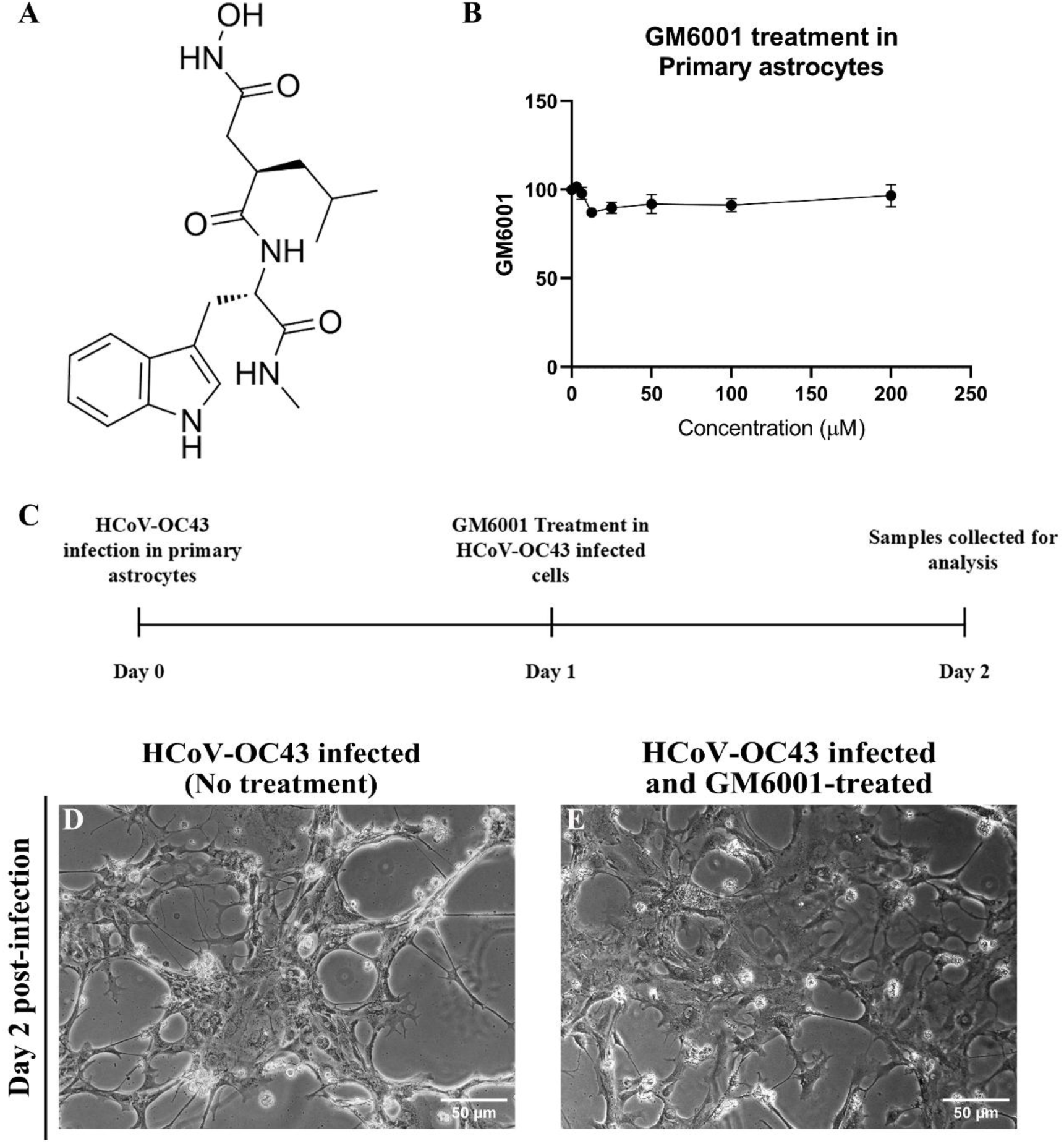
Determination of the working concentration of the broad-spectrum matrix metalloproteinase inhibitor GM6001 and experimental design for inhibitor treatment in HCoV-OC43-infected primary astrocytes. **(A)** Chemical structure of the broad-spectrum matrix metalloproteinase inhibitor GM6001 (adapted from MedChemExpress). **(B)** Primary astrocytes were treated with increasing concentrations of GM6001 (0-200 µM), and cell viability was assessed using the MTT assay. No significant reduction in MTT-based cell viability was observed across the concentrations tested. **(C)** Schematic representation of the experimental design showing HCoV-OC43 infection on Day 0, GM6001 treatment on Day 1, and sample collection on Day 2 post-infection. **(D-E)** Representative differential interference contrast (DIC) images of HCoV-OC43-infected astrocytes without treatment and following treatment with 25 µM GM6001, respectively, at Day 2 post-infection. No obvious treatment-associated cytotoxicity was observed following GM6001 treatment. Data represent the mean ± SEM from three independent experiments. Scale bar: 50 µm.

## GM6001 treatment restores junctional protein expression in HCoV-OC43-infected primary astrocytes

To determine whether MMP activity contributes to the junctional protein alterations observed following HCoV-OC43 infection, infected primary astrocytes were treated with GM6001, and the expression and localization of junctional proteins were examined. We first assessed the gap junction protein, Cx43. Untreated HCoV-OC43-infected astrocytes showed reduced punctate Cx43 staining at the cell periphery, whereas GM6001-treated infected astrocytes exhibited increased peripheral punctate Cx43 staining (Fig. 6A-B1, and white arrows in B1). The restoration of Cx43 localization was also observed in the additional panels of immunofluorescence images shown in Supplementary Figure 6. Immunoblot analysis further demonstrated increased Cx43 protein expression following GM6001 treatment, with densitometric analysis confirming a significant increase (Fig. 6C-D).

**Figure 6.**
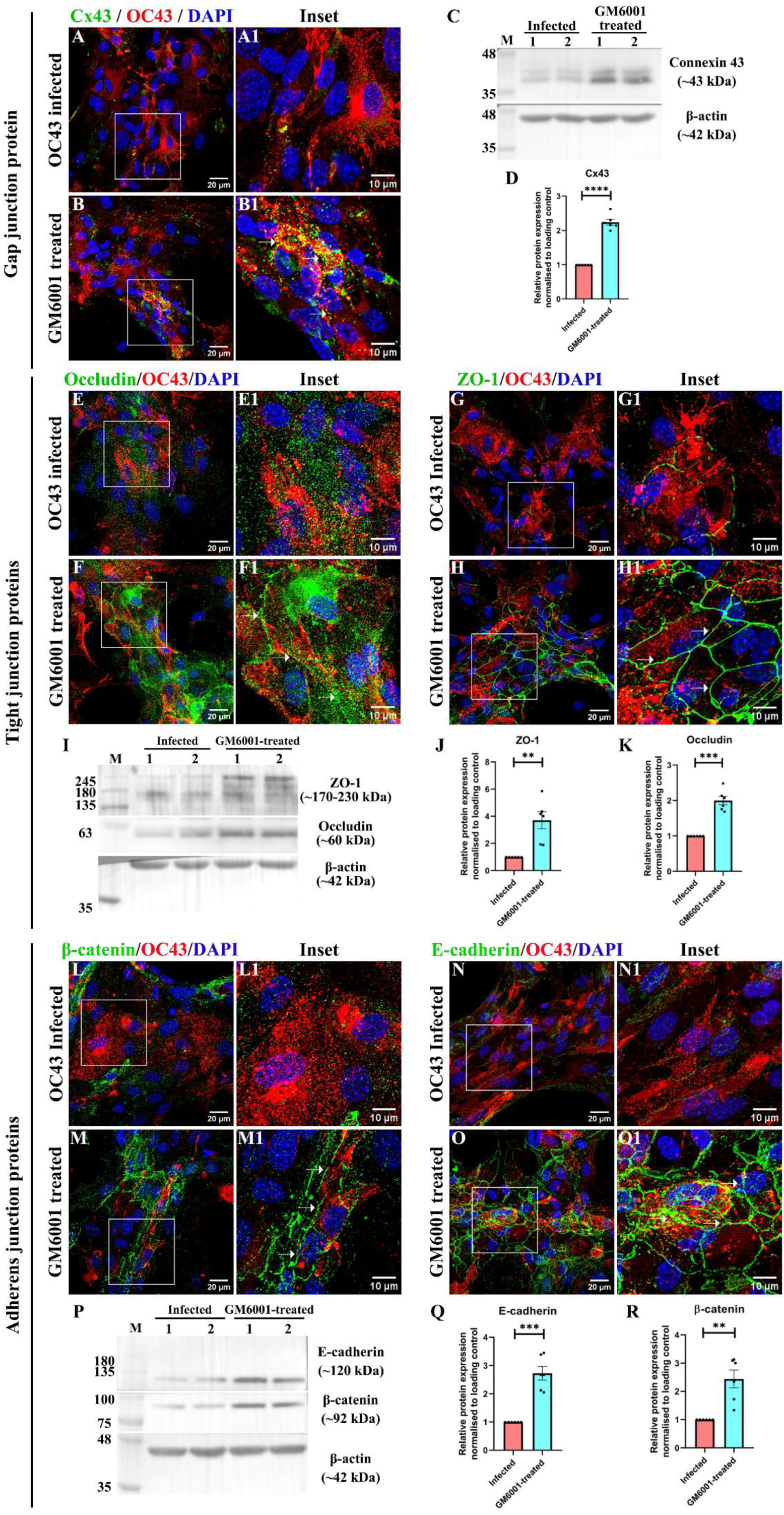
GM6001 treatment restores junctional protein expression in HCoV-OC43-infected primary astrocytes. Primary astrocytes infected with HCoV-OC43 were treated with 25 µM GM6001 and analyzed for junctional protein expression by double-label immunofluorescence and immunoblotting. (A-B) Double-label immunofluorescence staining showed Cx43 expression in untreated HCoV-OC43-infected astrocytes (A) and GM6001-treated infected astrocytes (B). GM6001-treated infected astrocytes showed increased peripheral Cx43 staining. (B1) The inset highlights restored Cx43 staining in GM6001-treated infected cells, with white arrows indicating representative regions. (C) Immunoblot analysis showed Cx43 and β-actin protein expression in untreated and GM6001-treated HCoV-OC43-infected astrocytes. (D) Densitometric quantification of Cx43 protein expression normalized to β-actin. (E-F) Double-label immunofluorescence staining showed Occludin expression in untreated HCoV-OC43-infected astrocytes (E) and GM6001-treated infected astrocytes (F). Untreated infected astrocytes exhibited disrupted Occludin localization, whereas GM6001-treated infected astrocytes showed increased membrane-associated staining. (F1) The inset highlights restored Occludin staining in GM6001-treated infected cells, with white arrows indicating representative regions. (G-H) Double-label immunofluorescence staining showed ZO-1 expression in untreated HCoV-OC43-infected astrocytes (G) and GM6001-treated infected astrocytes (H). Untreated infected astrocytes displayed disrupted ZO-1 localization, whereas GM6001-treated infected astrocytes showed increased membrane-associated staining. (H1) The inset highlights restored ZO-1 staining in GM6001-treated infected cells, with white arrows indicating representative regions. (I) Immunoblot analysis showed Occludin, ZO-1, and β-actin protein expression in untreated and GM6001-treated HCoV-OC43-infected astrocytes. (J-K) Densitometric quantification of ZO-1 and Occludin protein expression, respectively, normalized to β-actin. (L-M) Double-label immunofluorescence staining showed β-catenin expression in untreated HCoV-OC43-infected astrocytes (L) and GM6001-treated infected astrocytes (M). Untreated infected astrocytes exhibited reduced and fragmented β-catenin localization, whereas GM6001-treated infected astrocytes showed increased membrane-associated staining. (M1) The inset highlights restored β-catenin staining in GM6001-treated infected cells, with white arrows indicating representative regions. (N-O) Double-label immunofluorescence staining showed E-cadherin expression in untreated HCoV-OC43-infected astrocytes (N) and GM6001-treated infected astrocytes (O). Untreated infected astrocytes exhibited reduced and fragmented E-cadherin localization, whereas GM6001-treated infected astrocytes showed increased membrane-associated staining. (O1) The inset highlights restored E-cadherin staining in GM6001-treated infected cells, with white arrows indicating representative regions. (P) Immunoblot analysis showed E-cadherin, β-catenin, and β-actin protein expression in untreated and GM6001-treated HCoV-OC43-infected astrocytes. (Q-R) Densitometric quantification of E-cadherin and β-catenin protein expression, respectively, normalized to β-actin. Data represent mean ± SEM from three independent experiments. Statistical significance: *p < 0.05, **p < 0.01, ***p < 0.001, and ****p < 0.0001. Scale bars: 20 µm; inset scale bars: 10 µm.

We next examined the effect of GM6001 on tight junction proteins. Untreated HCoV-OC43-infected astrocytes displayed disrupted and diffuse Occludin and ZO-1 staining, whereas GM6001-treated infected astrocytes showed increased membrane-associated localization of both proteins (Fig. 6E-H1, and white arrows in F1 and H1). Similar restoration of Occludin and ZO-1 localization was observed in the additional panels of immunofluorescence images presented in Supplementary Figure 7. Immunoblot analysis showed increased protein expression of both Occludin and ZO-1 following GM6001 treatment, with densitometric analysis confirming these increases (Fig. 6I-K). Although GM6001 treatment improved Occludin abundance and its membrane-associated distribution, its localization did not fully resemble the junctional pattern observed in control astrocytes, suggesting that some aspects of Occludin trafficking or localization may remain impaired.

Finally, we examined the adherens junction proteins β-catenin and E-cadherin. Untreated HCoV-OC43-infected astrocytes exhibited reduced and fragmented membrane-associated staining of both proteins, whereas GM6001-treated infected astrocytes showed increased membrane-associated localization (Fig. 6L-O1, and white arrows in M1 and O1). The restoration of β-catenin and E-cadherin localization was further supported by the additional panels of immunofluorescence images shown in Supplementary Figure 8. Immunoblot analysis demonstrated increased E-cadherin and β-catenin protein expression following GM6001 treatment, as confirmed by densitometric analysis (Fig. 6P-R). Together, these findings indicate that inhibition of MMP activity with GM6001 attenuates the HCoV-OC43-associated reduction and redistribution of multiple junctional proteins in primary astrocytes.

## Discussion

Astrocytes are resident glial cells of the central nervous system that are increasingly recognized as active participants in innate antiviral immunity. Following viral sensing or infection, astrocytes can initiate innate immune signaling and produce type I interferons, cytokines, and chemokines, thereby contributing to the restriction of viral replication and coordinating responses with neighbouring cells, including microglia^19^. At the same time, sustained or dysregulated astrocyte responses can alter the cellular environment and contribute to neuroinflammation and tissue dysfunction. Importantly, viral infection can also directly alter the cellular architecture of astrocytes, including junctional proteins that are essential for intercellular communication and cellular organization. Such virus-induced remodeling of astrocytic junctions has been reported for several viruses, including SARS-CoV-2, Borna disease virus, influenza virus, and HIV ^8,20–22^.

Our previous studies using the neurotropic murine β-coronavirus MHV-A59 established that coronavirus infection alters astrocytic Cx43 expression, trafficking, and gap-junctional communication, both in primary astrocytes and in the infected CNS^9,23^ . Subsequent studies from our group demonstrated that MHV-A59-mediated disruption of Cx43 trafficking involves the microtubule network and that modulation of the ERp29-Cx43 axis can restore Cx43 trafficking and gap-junctional communication^10,13,14^ . These findings, together with growing evidence that astrocytes participate actively in antiviral immune responses, highlight the importance of understanding how viral infection simultaneously engages astrocytic immune signaling and alters cellular junctional organization.

In the present study, we used HCoV-OC43 infection of primary astrocytes to investigate whether coronavirus infection produces broader remodeling of astrocytic junctional proteins and to define the cellular mechanisms associated with these changes. Infection resulted in reduced expression and altered membrane localization of the gap junction protein Cx43, the tight junction proteins Occludin and ZO-1, and the adherens junction proteins β-catenin and E-cadherin. Mechanistically, infection was accompanied by Golgi apparatus disruption, an NF-κB-associated inflammatory response, and MMP9 upregulation, while ROS levels decreased and the ERK/MAPK pathway showed no evidence of activation. Importantly, pharmacological inhibition of MMP activity with GM6001 restored the expression and membrane-associated localization of all junctional proteins. Together, these findings identify MMP activity as a functional contributor to HCoV-OC43-induced junctional remodeling and provide a framework for understanding how coronavirus infection may alter astrocyte function and CNS homeostasis.

The simultaneous alteration of gap junction, tight junction, and adherens junction proteins is a central finding of this study. Previous work with the neurotropic murine β-coronavirus MHV-A59 has demonstrated selective loss of Cx43 in infected astrocytes, whereas the present study demonstrates a broader phenotype involving multiple junctional compartments. Gap junctions, tight junctions, and adherens junctions are supported by distinct but interconnected scaffolding and trafficking systems, and their simultaneous disruption suggests that HCoV-OC43 induces coordinated remodeling rather than targeting a single junctional component. The reduction of Cx43 may affect astrocytic intercellular communication, while changes in Occludin and ZO-1 may alter junction-associated organization at astrocytic processes. The concurrent reduction of β-catenin and E-cadherin is particularly relevant because the cadherin-catenin complex links cell-cell adhesion to the cortical actin cytoskeleton and contributes to the organization and polarity of astrocytic processes^24^. Moreover, adherens junctions, tight junctions, and gap junctions exhibit molecular and functional crosstalk, mediated in part by shared scaffolding proteins such as ZO-1 and interactions between Cx43 and junction-associated proteins^25,26^. Thus, disruption of one junctional compartment may influence the stability of the others, providing a plausible basis for the broad junctional phenotype observed following HCoV-OC43 infection.

The observed junctional remodeling was accompanied by disruption of the Golgi apparatus, providing a potential mechanism for impaired junctional protein homeostasis. Junctional proteins undergo biosynthetic processing and intracellular trafficking before reaching the plasma membrane. The Golgi apparatus contributes to the processing and delivery of several junctional proteins, including Cx43 and E-cadherin, and disruption of this trafficking pathway may therefore contribute to altered membrane-associated abundance of junctional proteins^27,28^ . Coronavirus-induced Golgi fragmentation has also been reported in other cellular systems, and our previous work demonstrated that HCoV-OC43 infection can impair Golgi-associated Cx43 trafficking in A549 cells^12^ . The present findings extend these observations to primary astrocytes and, importantly, show that Golgi disruption occurs together with altered localization of multiple junctional proteins rather than Cx43 alone.

A second mechanistic component identified in this study was an NF-κB-associated inflammatory response. HCoV-OC43 infection was associated with increased expression of p65/RelA and IKKβ at the protein level, together with increased p65/RelA transcripts, supporting an NF-κB-associated inflammatory response. This response was accompanied by induction of inflammatory cytokines and chemokines, including CCL5, IL-6, TNF-α, and IL-1β, as well as type I interferon responses involving IFN-α and IFN-β, while IFN-γ remained unchanged. At the same time, increased IL-10 and TGF-β suggest the presence of counter-regulatory signals that may limit excessive inflammation. The induction of Nfkbia, which encodes IκBα, further indicates a potential negative-feedback mechanism within the NF-κB regulatory network. Although the current experiments do not establish direct NF-κB activation through nuclear translocation or phosphorylation, the coordinated increase in NF-κB pathway-associated components and inflammatory mediators identifies this pathway as a plausible upstream contributor to the MMP-associated response. Pharmacological inhibition of NF-κB with BAY 11-7082 was also explored. However, concentrations tolerated by control astrocytes exhibited substantial loss of cell viability in HCoV-OC43-infected primary astrocytes. Hence, direct causal linkage between NF-κB and MMP9 induction will require further investigation with another NF-κB inhibitor.

The analysis of potential upstream pathways further refined the mechanistic model. Unexpectedly, HCoV-OC43 infection resulted in reduced cellular ROS levels, and Erk2 protein expression was significantly downregulated. Thus, the present data do not support ROS/ERK-MAPK signaling as a major driver of the MMP-associated response under the experimental conditions used. This observation is notable because ROS generation and ERK/MAPK signaling are frequently implicated in MMP9 regulation in other inflammatory and disease models^29–31^ . The reduced ROS response therefore suggests that HCoV-OC43-induced MMP-associated junctional remodeling in astrocytes may proceed through alternative signaling mechanisms.

Given the concurrent NF-κB-associated inflammatory response and the induction of MMP9, we next examined whether MMP activity contributes functionally to the junctional phenotype. HCoV-OC43 infection significantly increased MMP9 transcript levels, while treatment with the broad-spectrum MMP inhibitor GM6001 restored the expression and membrane-associated localization of Cx43, Occludin, ZO-1, β-catenin, and E-cadherin. This pharmacological rescue provides functional evidence that MMP activity contributes to the junctional remodeling observed during infection. MMP9 is a biologically plausible candidate because MMP-mediated cleavage of junctional and barrier-associated proteins has been reported in inflammatory, BBB, and viral infection models, and astrocytic MMP9 has been associated with junctional and barrier injury^32–34^ .

GM6001 was selected as a broad-spectrum MMP inhibitor because the objective of this experiment was initially to determine whether inhibition of MMP activity as a group could attenuate the junctional remodeling phenotype, rather than to assign the effect specifically to MMP9. The restoration of multiple junctional proteins following broad-spectrum MMP inhibition therefore provides functional evidence for the involvement of MMP activity while avoiding an overinterpretation of MMP9 as the sole protease responsible for the phenotype. Therefore, MMP activity, with MMP9 representing a prominent candidate, contributes to HCoV-OC43-induced junctional remodeling. The restoration of multiple junctional proteins following MMP inhibition indicates that extracellular proteolytic activity is functionally relevant to the observed phenotype.

Taken together, the findings support a model in which HCoV-OC43 infection induces two mechanistically distinct but potentially converging processes: Golgi apparatus disruption and an NF-κB-associated inflammatory response leading to MMP9 induction. Golgi dysfunction may compromise the trafficking of junctional proteins, whereas MMP activity may promote the loss of membrane-associated junctional proteins through extracellular proteolysis. The convergence of these processes could explain why proteins belonging to gap junctions, tight junctions, and adherens junctions are simultaneously affected. The accompanying inflammatory response, including type I interferons and pro-inflammatory cytokines, together with compensatory IL-10, TGF-β, and IκBα responses, further highlights the ability of astrocytes to act both as mediators and regulators of antiviral inflammation (Fig. 7).

**Figure 7:**
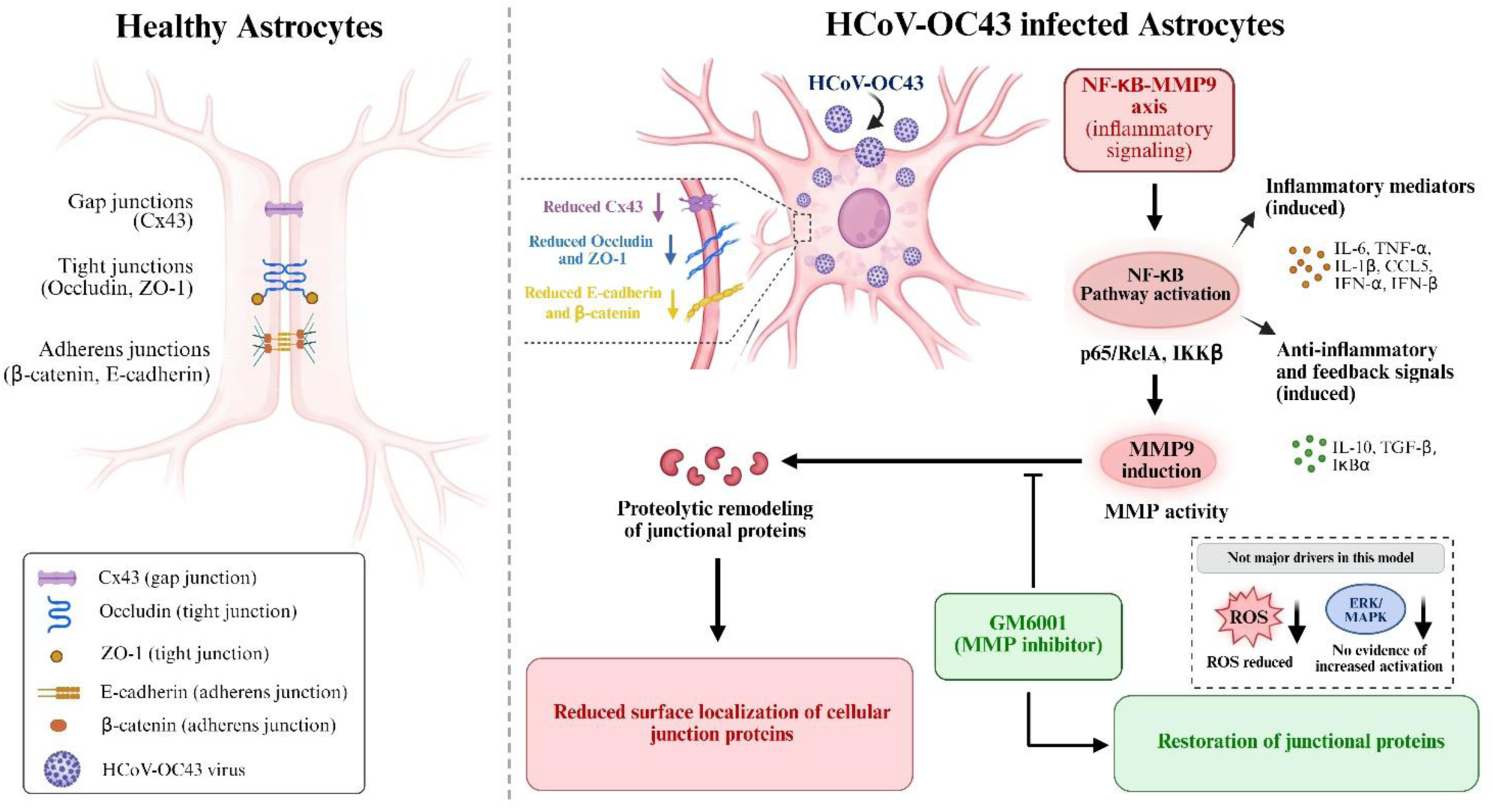
HCoV-OC43 infection is associated with astrocytic junctional remodeling and an inflammatory response. Infection of primary astrocytes with human coronavirus OC43 (HCoV-OC43) alters the expression and localization of gap junction, tight junction, and adherens junction proteins, including Connexin 43 (Cx43), Occludin, ZO-1, β-catenin, and E-cadherin. Infection is also associated with increased expression of NF-κB pathway-associated components and inflammatory mediators, together with increased MMP9 transcript levels. Pharmacological inhibition of matrix metalloproteinase (MMP) activity with GM6001 improves junctional protein abundance and membrane localization, supporting a role for MMP activity in junctional remodeling.

These findings have potential physiological relevance because astrocytes are integral components of the neurovascular unit and contribute to CNS homeostasis through their endfeet, intercellular communication, and interactions with neighboring cells and vascular structures^35^. Alteration of Cx43 could affect astrocytic gap-junction communication, while loss of Occludin and ZO-1 may alter junction-associated organization at astrocytic processes. Similarly, disruption of the E-cadherin/β-catenin complex may affect cell-cell adhesion and cytoskeletal organization. Previous studies of MHV-A59 infection have demonstrated functional impairment of astrocytic gap-junction communication, providing biological support for the possibility that the Cx43 alterations observed here have functional consequences^9,10,13,14^ . However, because the present study did not directly measure dye coupling, barrier permeability, or endfoot polarization, the functional consequences of the observed junctional remodeling remain to be established. Importantly, these findings should not be interpreted as direct evidence of BBB disruption, since endothelial barrier function was not assessed.

The broader relevance of this mechanism is supported by observations that other human β-coronaviruses, including SARS-CoV, SARS-CoV-2, and MERS-CoV, can alter junctional proteins and barrier-associated structures in epithelial and endothelial systems^8,36–39^. SARS-CoV-2 infection has also been associated with BBB dysfunction and increased MMP9 in experimental models, although the specific junctional proteins affected can vary according to cell type and model^40^. Thus, the present findings raise the possibility that coronavirus-induced junctional remodeling may represent a broader feature of coronavirus-associated cellular dysfunction. HCoV-OC43 provides a useful BSL-2 model in which such mechanisms can be investigated experimentally and may therefore help define cellular processes relevant to more pathogenic β-coronaviruses.

The possible relevance to persistent neurological consequences, including those reported after SARS-CoV-2 infection, should nevertheless be considered cautiously. The present study examines acute HCoV-OC43 infection in primary murine astrocytes and does not address viral persistence, recovery after viral clearance, or post-acute neurological disease. Our previous MHV-A59 studies showed that junctional proteins can recover differentially following viral clearance, raising the possibility that the persistence or reversibility of junctional remodeling may be an important determinant of long-term consequences^9^. Whether HCoV-OC43-induced junctional changes persist after viral clearance therefore remains an important question for future studies.

The therapeutic implications of the GM6001 rescue are also potentially important but require further validation. Restoration of multiple junctional proteins following broad-spectrum MMP inhibition provides proof of concept that targeting extracellular proteolytic activity may help preserve junctional integrity during coronavirus infection. However, the broad activity of GM6001 limits its interpretation and therapeutic specificity. Selective MMP9 inhibition, genetic depletion of MMP9, direct assessment of MMP9 enzymatic activity, and evaluation of other MMP family members will be required to determine the specific protease(s) responsible.

Several limitations should be considered when interpreting these findings. The study was performed in primary astrocytes isolated from neonatal C57BL/6 mice, and therefore the relevance of the observed mechanisms to human astrocytes and the intact CNS requires validation in human astrocytes and appropriate neurovascular models. GM6001 is broad-spectrum and cannot establish MMP9 as the specific mediator, while MMP9 was measured at the transcript level without direct assessment of enzymatic activity. The study also did not directly measure BBB permeability, functional gap-junction communication, or astrocyte endfoot organization, and therefore the physiological consequences of junctional remodeling remain inferential. In addition, the upstream mechanism connecting HCoV-OC43 infection to the NF-κB-associated response remains unresolved, and the unexpected reduction in ROS and lack of evidence for increased ERK/MAPK signaling warrant further investigation. Future studies using human astrocytes, neurovascular unit models, selective or genetic MMP9 approaches, direct NF-κB activation assays, and extended infection and recovery time courses will be important to establish the physiological and mechanistic significance of these observations.

Overall, this study identifies broad remodeling of astrocytic junctional architecture as an important cellular response to HCoV-OC43 infection. The convergence of Golgi disruption and MMP-associated activity, with the NF-κB-associated inflammatory response providing a potential upstream context for MMP9 induction, provides a mechanistic framework linking coronavirus infection to alterations in multiple junctional compartments. These findings extend the understanding of how β-coronavirus infection may perturb astrocyte homeostasis and provide a basis for investigating whether preservation of junctional integrity can mitigate coronavirus-associated CNS dysfunction.

## Acknowledgments

The authors sincerely thank the Department of Biological Sciences, Indian Institute of Science Education and Research (IISER) Kolkata, and the IISER Kolkata Animal Facility for providing essential infrastructure and support. We thank Dr. Sourish Ghosh (Indian Institute of Chemical Biology, Kolkata, India) for kindly providing Vero cells and Dr. Brenda Hogue (Arizona State University, USA) for generously providing the anti-OC43 antibody. We gratefully acknowledge BEI Resources, NIAID, NIH, for providing the HCoV-OC43 virus. We also acknowledge the IISER Kolkata Central Imaging Facility and Mr. Ritabrata Ghosh for technical assistance. The authors thank Mr. Bishal Hazra for assistance with flow cytometry data acquisition. The authors thank Prof. Michael Koval (Emory University School of Medicine, USA) for critically reviewing the final draft of the manuscript. S.K. acknowledges the Council of Scientific and Industrial Research (CSIR), India, for fellowship support. The authors also sincerely acknowledge the funding agencies listed in the Funding section for their support.

## Author Contributions

**S.K.**: Conceptualization, methodology, software, validation, formal analysis, investigation, resources, data curation, writing - original draft preparation, writing - review and editing, visualization.

**J.D.S.**: Conceptualization, methodology, validation, formal analysis, investigation, resources, writing - review and editing, visualization, supervision.

Both authors have read and agreed to the published version of the manuscript.

## References

1. Morgello, S. (2020). Coronaviruses and the central nervous system. J. Neurovirol. 26, 459–473. 10.1007/s13365-020-00868-7.

2. Iroegbu, J.D., Ifenatuoha, C.W., and Ijomone, O.M. (2020). Potential neurological impact of coronaviruses: implications for the novel SARS-CoV-2. Neurological Sciences 41, 1329–1337. 10.1007/s10072-020-04469-4.

3. Lawrence, J.M., Schardien, K., Wigdahl, B., and Nonnemacher, M.R. (2023). Roles of neuropathology-associated reactive astrocytes: a systematic review. Acta Neuropathol. Commun. 11, 42. 10.1186/s40478-023-01526-9.

4. Wu, D., Chen, Q., Chen, X., Han, F., Chen, Z., and Wang, Y. (2023). The blood–brain barrier: structure, regulation, and drug delivery. Preprint at Springer Nature, 10.1038/s41392-023-01481-w.

5. Jiang, H., Zhang, Y., Wang, Z.-Z., and Chen, N.-H. (2023). Connexin 43: An Interface Connecting Neuroinflammation to Depression. Molecules 28, 1820. 10.3390/molecules28041820.

6. Boardman, S.A., Hetherington, C., Hughes, T., Cook, C., Galea, I., Hilton, O., Solomon, T., Luster, A.D., Allan, S., Kurt-Jones, E., et al. (2025). Viral Infection and the Blood-Brain Barrier: Molecular Research Insights and Therapies. J. Infect. Dis. 10.1093/infdis/jiaf455.

7. Krasemann, S., Haferkamp, U., Pfefferle, S., Woo, M.S., Heinrich, F., Schweizer, M., Appelt-Menzel, A., Cubukova, A., Barenberg, J., Leu, J., et al. (2022). The blood-brain barrier is dysregulated in COVID-19 and serves as a CNS entry route for SARS-CoV-2. Stem Cell Reports 17, 307–320. 10.1016/j.stemcr.2021.12.011.

8. Reynolds, J.L., and Mahajan, S.D. (2021). SARS-COV2 Alters Blood Brain Barrier Integrity Contributing to Neuro-Inflammation. Preprint at Springer, 10.1007/s11481-020-09975-y.

9. Basu, R., Banerjee, K., Bose, A., and Das Sarma, J. (2016). Mouse Hepatitis Virus Infection Remodels Connexin43-Mediated Gap Junction Intercellular Communication In Vitro and In Vivo . J. Virol. 90, 2586–2599. 10.1128/jvi.02420-15.

10. Basu, R., Bose, A., Thomas, D., and Sarma, J. Das (2017). Microtubule-assisted altered trafficking of astrocytic gap junction protein connexin 43 is associated with depletion of connexin 47 during mouse hepatitis virus infection. Journal of Biological Chemistry 292, 14747–14763. 10.1074/jbc.M117.786491.

11. Tress, O., Maglione, M., May, D., Pivneva, T., Richter, N., Seyfarth, J., Binder, S., Zlomuzica, A., Seifert, G., Theis, M., et al. (2012). Panglial Gap Junctional Communication is Essential for Maintenance of Myelin in the CNS. The Journal of Neuroscience 32, 7499–7518. 10.1523/JNEUROSCI.0392-12.2012.

12. Karmakar, S., and Das Sarma, J. (2024). Human coronavirus OC43 infection remodels connexin 43-mediated gap junction intercellular communication in vitro . J. Virol. 98. 10.1128/jvi.00478-24.

13. Bose, A., Kasle, G., Jana, R., Maulik, M., Thomas, D., Mulchandani, V., Mukherjee, P., Koval, M., and Das Sarma, J. (2023). Regulatory role of endoplasmic reticulum resident chaperone protein ERp29 in anti-murine β-coronavirus host cell response. Journal of Biological Chemistry 299, 102836. 10.1016/j.jbc.2022.102836.

14. Kasle, G., Sharma, M., Kumar, S., Das, P., Das Sarma, S., Koval, M., Kenyon, L.C., and Das Sarma, J. (2025). Phenylbutyrate restricts murine β-coronavirus infectivity and limits virus-induced demyelination in vivo. J Virol 99, e0081925. 10.1128/jvi.00819-25.

15. Samal, S.K., Sharma, M., and Sarma, J. Das (2024). Isolation and Enrichment of Major Primary Neuroglial Cells from Neonatal Mouse Brain. Bio. Protoc. 14. 10.21769/BioProtoc.4921.

16. Marek, R., Caruso, M., Rostami, A., Grinspan, J.B., and Sarma, J. Das (2008). Magnetic cell sorting: A fast and effective method of concurrent isolation of high purity viable astrocytes and microglia from neonatal mouse brain tissue. J. Neurosci. Methods 175, 108–118. 10.1016/j.jneumeth.2008.08.016.

17. Mondal, A., Das, B., Karmakar, S., Pani, S., Khan, S., Gupta, P., and Das Sarma, J. (2024). Modulatory Role of Pantropic Cell Signaling Pathways in the Antimigratory and Antiproliferative Action of Triazole Chelated Iridium(III) Complexes in Cervical Cancer Cells. J. Med. Chem. 10.1021/acs.jmedchem.4c01997.

18. Banerjee, S., Paul, S., Majumder, R., Manna, D., Mondal, S., Dhar, D., Karmakar, S., Sarma, J. das, Das, A., Singh, N.D.P., et al. (2026). Coupling DNA intercalation with redox catalysis: selective killing mechanism of glioblastoma by daidzin. Nucleic Acids Res. 54. 10.1093/nar/gkag397.

19. Daniels, B.P., Jujjavarapu, H., Durrant, D.M., Williams, J.L., Green, R.R., White, J.P., Lazear, H.M., Gale, M., Diamond, M.S., and Klein, R.S. (2017). Regional astrocyte IFN signaling restricts pathogenesis during neurotropic viral infection. Journal of Clinical Investigation 127, 843–856. 10.1172/JCI88720.

20. Köster-Patzlaff, C., Hosseini, S.M., and Reuss, B. (2007). Persistent Borna Disease Virus infection changes expression and function of astroglial gap junctions in vivo and in vitro. Brain Res. 1184, 316–332. 10.1016/j.brainres.2007.09.062.

21. Ruan, T., Sun, J., Liu, W., Prinz, R.A., Peng, D., Liu, X., and Xu, X. (2020). H1N1 Influenza Virus Cross-Activates Gli1 to Disrupt the Intercellular Junctions of Alveolar Epithelial Cells. Cell Rep. 31. 10.1016/j.celrep.2020.107801.

22. Sufiawati, I., and Tugizov, S.M. (2014). HIV-Associated Disruption of Tight and Adherens Junctions of Oral Epithelial Cells Facilitates HSV-1 Infection and Spread. PLoS One 9, e88803. 10.1371/journal.pone.0088803.

23. Bose, A., Basu, R., Maulik, M., and Das Sarma, J. (2018). Loss of Cx43-Mediated Functional Gap Junction Communication in Meningeal Fibroblasts Following Mouse Hepatitis Virus Infection. Mol. Neurobiol. 55, 6558–6571. 10.1007/s12035-017-0861-3.

24. Arikkath, J., and Reichardt, L.F. (2008). Cadherins and catenins at synapses: roles in synaptogenesis and synaptic plasticity. Trends Neurosci. 31, 487–494. 10.1016/j.tins.2008.07.001.

25. Strauss, R.E., and Gourdie, R.G. (2020). Cx43 and the Actin Cytoskeleton: Novel Roles and Implications for Cell-Cell Junction-Based Barrier Function Regulation. Biomolecules 10, 1656. 10.3390/biom10121656.

26. Giepmans, B. (2004). Gap junctions and connexin-interacting proteins. Cardiovasc. Res. 62, 233–245. 10.1016/j.cardiores.2003.12.009.

27. Laird, D.W. (2006). Life cycle of connexins in health and disease. Biochemical Journal 394, 527–543. 10.1042/BJ20051922.

28. Harris, T.J.C., and Tepass, U. (2010). Adherens junctions: from molecules to morphogenesis. Nat. Rev. Mol. Cell Biol. 11, 502–514. 10.1038/nrm2927.

29. Lian, S., Xia, Y., Khoi, P.N., Ung, T.T., Yoon, H.J., Kim, N.H., Kim, K.K., and Jung, Y. Do (2015). Cadmium induces matrix metalloproteinase-9 expression via ROS-dependent EGFR, NF-κB, and AP-1 pathways in human endothelial cells. Toxicology 338, 104–116. 10.1016/j.tox.2015.10.008.

30. Al-Sadi, R., Engers, J., Haque, M., King, S., Al-Omari, D., and Ma, T.Y. (2021). Matrix Metalloproteinase-9 (MMP-9) induced disruption of intestinal epithelial tight junction barrier is mediated by NF-κB activation. PLoS One 16. 10.1371/journal.pone.0249544.

31. Ranaivo, H.R., Hodge, J.N., Choi, N., and Wainwright, M.S. (2012). Albumin induces upregulation of matrix metalloproteinase-9 in astrocytes via MAPK and reactive oxygen species-dependent pathways. J. Neuroinflammation 9. 10.1186/1742-2094-9-68.

32. Vermeer, P.D., Denker, J., Estin, M., Moninger, T.O., Keshavjee, S., Karp, P., Kline, J.N., and Zabner, J. (2009). MMP9 modulates tight junction integrity and cell viability in human airway epithelia. Am J Physiol Lung Cell Mol Physiol 296, 751–762. 10.1152/ajplung.90578.2008.-The.

33. Yang, F., Zhao, K., Zhang, X., Zhang, J., and Xu, B. (2016). ATP Induces Disruption of Tight Junction Proteins via IL-1 Beta-Dependent MMP-9 Activation of Human Blood-Brain Barrier *In Vitro*. Neural Plast. 2016, 1–12. 10.1155/2016/8928530.

34. Min, H., Hong, J., Cho, I.-H., Jang, Y.H., Lee, H., Kim, D., Yu, S.-W., Lee, S., and Lee, S.J. (2015). TLR2-induced astrocyte MMP9 activation compromises the blood brain barrier and exacerbates intracerebral hemorrhage in animal models. Mol. Brain 8, 23. 10.1186/s13041-015-0116-z.

35. Abbott, N.J., Rönnbäck, L., and Hansson, E. (2006). Astrocyte–endothelial interactions at the blood–brain barrier. Nat. Rev. Neurosci. 7, 41–53. 10.1038/nrn1824.

36. Raghavan, S., Kenchappa, D.B., and Leo, M.D. (2021). SARS-CoV-2 Spike Protein Induces Degradation of Junctional Proteins That Maintain Endothelial Barrier Integrity. Front. Cardiovasc. Med. 8. 10.3389/fcvm.2021.687783.

37. Alvarez, F., de Melo, G.D., Larrous, F., Kergoat, L., Boëda, B., Michel, V., Seilhean, D., Tichit, M., Hing, D., Hardy, D., et al. (2025). The SARS-CoV-2 envelope PDZ binding motif acts as a virulence factor disrupting host’s epithelial cell–cell junctions. Cell. Mol. Biol. Lett. 30. 10.1186/s11658-025-00758-y.

38. Teoh, K.-T., Siu, Y.-L., Chan, W.-L., Schlüter, M.A., Liu, C.-J., Peiris, J.S.M., Bruzzone, R., Margolis, B., and Nal, B. (2010). The SARS Coronavirus E Protein Interacts with PALS1 and Alters Tight Junction Formation and Epithelial Morphogenesis. Mol. Biol. Cell 21, 3838–3852. 10.1091/mbc.e10-04-0338.

39. Chiok, K.L., Jenik, K., Fenton, M., Falzarano, D., Dhar, N., and Banerjee, A. (2025). MERS-CoV and SARS-CoV-2 infection in diverse human lung organoid-derived cultures. J. Virol. 99. 10.1128/jvi.01098-25.

40. Kempuraj, D., Tsilioni, I., Aenlle, K.K., Klimas, N.G., and Theoharides, T.C. (2024). Long COVID elevated MMP-9 and release from microglia by SARS-CoV-2 Spike protein. Transl. Neurosci. 15. 10.1515/tnsci-2022-0352.

